# Evolution and Genetic Diversity of the Retroviral Envelope in Anamniotes

**DOI:** 10.1101/2021.11.23.469661

**Authors:** Yicong Chen, Xiaojing Wang, Meng-En Liao, Yuhe Song, Yu-Yi Zhang, Jie Cui

## Abstract

Retroviruses are widely distributed in all vertebrates, as are their endogenous form, endogenous retrovirus (ERV), which serves as “fossil” evidence to trace the ancient origins and history of virus-host interactions over millions of years. The retroviral envelope (Env) plays a significant role in host range determination, but major information on genetic diversification in anamniotes is lacking. Here, by incorporating multiple-round *in silico* similarity search and phylogenomic analysis, 25498 ERVs with gamma-type Env (GTE), covalently associated Env, were discovered by searching against all 974 available fish and 19 amphibian genomes, but no beta-type Env (BTE), noncovalently associated Env, were found. Furthermore, a nine-type classification system of anamniote GTE was proposed by combining phylogenetic and domain/motif analyses. The elastic genomic organization and overall phylogenetic incongruence between anamniotic Env and its neighboring polymerase (Pol) implied that early retroviral diversification in anamniotic vertebrates was facilitated by frequent recombination. At last, host opioid growth factor receptor (OGFr) gene capturing by anamniotic ERVs with GTE was reported for the first time. Overall, our findings overturn traditional Pol genotyping and reveal a complex evolutionary history of anamniotic retroviruses inferred by Env evolution.

**Author summary:** Although the retroviral envelope (Env) protein in amniotes has been well studied, its evolutionary history in anamniotic vertebrates is ambiguous. By analyzing more than 25000 ERVs with gamma-type Env (GTE) in anamniotes, several important evolutionary features were identified. First, GTE were found to be widely distributed among different amphibians and fish. Second, nine types of GTE were discovered, revealing the great genetic diversity. Third, GTE-containing ERVs have rampantly proliferated in certain amphibians such as *Ambystoma mexicanum*, and the copy number was found to be markedly higher than 10000. Fourth, the incongruence between the Env and Pol phylogenies suggested that frequent recombination shaped the early evolution of anamniote retroviruses. Fifth, an ancient horizontal gene transfer event was discovered from anamniotes to ERVs with GTE. These findings reveal a complex evolution pattern for retroviral Env in anamniotes.

## Introduction

Retroviruses (RVs) (family Retroviridae), which were first discovered more than a hundred years ago [1], are medically and economically important because some are associated with severe infectious diseases, cancer and immunodeficiency [2–4]. RVs occasionally integrate into the germline of the host and become endogenous retroviruses (ERVs), which serve as historical genomic ‘fossils’ for investigating viral origins and the history of virus-host interactions over millions of years [5, 6]. Due to the increasing number of sequenced vertebrate genomes, numerous ERVs have been uncovered, and some of these ERVs are distinct from the current exogenous RVs, as reflected by their genomic structure and genomic similarity, which indicates the ancient origin and long- term evolution of RVs [5].

Typical RVs contain three major protein-coding genes: group-specific antigen (gag), polymerase (pol) and envelope (env) genes. For the phylogenetic reconstruction of retroviral lineages, well-conserved Pol protein sequences, particularly the reverse transcriptase (RT) domain, are typically used, which allows the alignment of multiple ERVs or exogenous RVs from various vertebrates and can thus be used to infer the deep evolutionary history of RVs as a whole [7]. Based on the RT phylogeny, ERVs can be classified into three broad classes: 1) class I, which related to epsilonretroviruses and gammaretroviruses (GVs); 2) class II, which related to deltaretroviruses, lentiretroviruses, betaretroviruses and alpharetroviruses (AVs); and 3) class III, which related to spumaretroviruses.[8] Although an analysis based solely on Pol or RT has a distinct advantage, it also has a clear disadvantage in that it ignores many fine distinctions in other regions (such as env) of genomes among viruses [9–11].

Env, the encoded protein of which mediates entry into the host cell and thus determines the host range [11, 12], exhibits markedly higher variability than pol. This gene encodes two subunits: the surface subunit (SU) and the transmembrane fusion subunit (TM). While SU is the most variable region of the genome, TM is relatively well conserved across many RVs [11, 13, 14]. Env can be divided into two major types: covalently associated (gamma-type Env, GTE) and noncovalently associated (beta-type Env, BTE) [13]. These types can be readily distinguished by sequence similarity and the presence or absence of two regions: the immunosuppressive domain (ISD) and the CX6CC motif. A GTE variant from an alpharetrovirus (GTE-AV), which has an internal fusion peptide flanked by a pair of cysteines, has also been well characterized [11]. BTEs are typically harbored by betaretroviruses and lentiviruses and can only be found in mammals [13]. In contrast, GTEs are harbored by gammaretroviruses, deltaretroviruses, alpharetroviruses and betaretroviruses (recombinant RVs, including MPMV and SRV) and can be found in all five major classes of vertebrates (fishes, amphibians, reptiles, birds, and mammals) [9-11, 15, 16].

Although a substantial number of RVs-GTE (retroviruses with a gamma-type Env) have been identified across the evolutionary history of vertebrates [15, 17–19], these retroviruses in anamniote vertebrates remain poorly understood. Hence, in this study, by incorporating genomic and transcriptomic data and performing comprehensive phylogenomic analyses, we were able to identify hundreds of RV-GTE lineages, which significantly increased the known set of RVs-GTE in anamniote vertebrates. Most importantly, by combining phylogenetic and genomic structure characterization, we uncovered the structural complexity and phylogenetic diversity of GTEs and redefined the classification of Env. Additionally, some unique evolutionary features were revealed, which significantly increased our understanding of the diversity and evolution of RVs.

## Results

### Identification of ERVs-GTE, ERVs-BTE, expressed RVs-GTE and expressed RVs-BTE in amphibians and fish

To systematically identify ERVs-GTE (endogenous retroviruses with gamma-type Env) and ERVs-BTE (endogenous retroviruses with beta-type Env), all 974 available fish and 19 amphibian genomes (S1 Table) were screened using a combined multiple-round similarity searching approach (Fig 1). First, host genomes were screened to detect ERVs-GTE and ERVs-BTE using tBLASTn, and all Env sequences of representative RVs-GTE and RVs-BTE (S2 Table) were used as queries. Second, the significant hits were concatenated based on the host genomic location and alignment positions with reference RV proteins. These concatenated proteins were then further confirmed using phylogeny, and only sequences that clustered with RVs-GTE or RVs-BTE were included. Third, the same genomes were subjected to a second round of screening by tBLASTn using the confirmed concatenated Env protein sequences. Fourth, the significant hits that were also confirmed by phylogeny were extended to identify the full-length genome and classified into different lineages based on sequence similarity. Finally, the full-length viral genomes of each lineage were used as queries in a BLASTn search against the genome to determine the copies of each lineage (detailed in “Materials and Methods”). This analysis led to the discovery of 37,552 ERVs, with 25498 containing GTE in 37 vertebrate species, including 13 amphibians, 18 ray-finned fish and 6 cartilaginous fish, and no ERVs-BTE was found, which was consistent with the results of previous research showing that noncovalently associated Env (beta-type Env) could only be found in mammals [13] (S1 Data). These retroviral sequences were named in accordance with a previous nomenclature proposed for ERVs [8].

**Fig 1.**
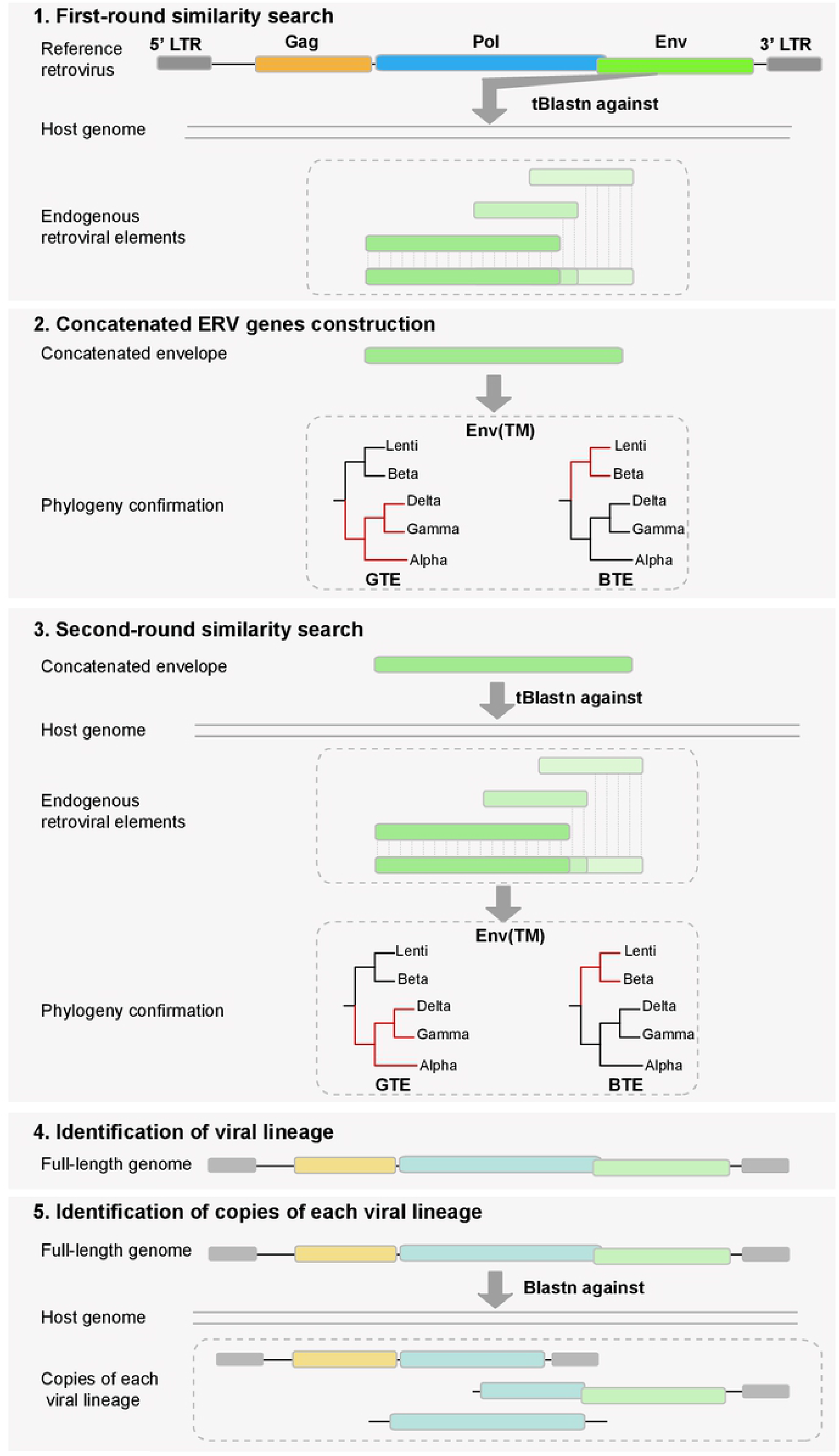
Schematic diagram of the pipeline to identify ERVs-GTE and ERVs-BTE. First, host genomes were screened to detect ERVs-GTE and ERVs-BTE using tBLASTn and all reference GTE and BTE sequences were used as queries. Second, the significant hits were concatenated based on the host genomic location and alignment positions with reference proteins. Then, these concatenated proteins were further confirmed using phylogeny and only sequences that clustered with GTE or BTE were included. Third, the same genomes were subjected to a second round of screening by tBLASTn using the confirmed concatenated protein sequences of Env. Fourth, the significant hits which were also confirmed by phylogeny were extended to identify the full-length genome and classified into different lineages based on sequence similarity. Finally, the full-length viral genomes of each lineage were used as queries to BLASTn search against genome to determine the copies of each lineage.

To better understand the expression of potential RVs-GTE, we also searched all available 378 fish and 61 amphibian RNA-seq datasets in the transcriptome sequence assembly (TSA) database (S1 Table). Notably, we found 1,093 retroviral Env contigs in 155 vertebrate species, including 29 amphibians, 113 ray-finned fish and 3 cartilaginous fish (S1 Data). These viral sequences were named in accordance with a previous nomenclature proposed for expressed RVs (exRVs) [20]. However, 143 of 155 species harboring such exRVs did not have any genomic data support. We still found that 5 amphibians (*Rana catesbeiana*, *Rhinatrema bivittatum*, *Microcaecilia unicolor*, *Pyxicephalus adspersus*, and *Rhinella marina*) and 7 ray-finned fish (*Astyanax mexicanus*, *Larimichthys crocea*, *Oncorhynchus kisutch*, *O. tshawytscha*, *Salmo trutta*, *Salvelinus alpinus*, and *Seriola dumerili*) harbored both endogenous and expressed forms of retroviral elements, and some of these shared high similarity (>95%) among the DNA and RNA copies in each host, which indicated the potential ability to express such viral elements. In addition, we found 4 copies of Env RNA contigs in *R. catesbeiana*, 1 copy in *P. adspersus*, 10 copies in *A. mexicanus*, 2 copies in *O. tshawytscha*, 2 copies in *S. trutta*, and 10 copies in *S. alpinus*; these copies were distantly related to the DNA copies that shared less than 50% similarity with each other, which implied the potential existence of exogenous forms of such viruses because most of these (21/29) encoded intact open reading frames (ORFs).

### Distribution of ERVs-GTE in amphibians and fish

The distribution of ERVs-GTE in different genomes was unbalanced. Ten of the 37 vertebrates, including 6 amphibians, 1 ray-finned fish and 3 cartilaginous fish, harbored more than 500 copies of ERVs-GTE, whereas 14 of the 37 vertebrates, including 3 amphibians, 9 ray-finned fish and 2 cartilaginous fish, harbored less than 20 copies (Fig 2A). Amphibians harbored more copies of ERVs-GTE than fish; in particular, *A. mexicanum* and *L. leishanense* had 20,943 and 7,517 copies, respectively. Surprisingly, the ERVs-GTE in *L. leishanense* constituted 1.27% of the genome, indicating rampant proliferation and good adaptation of such viral elements to their toad host. In addition, we found that ray-finned fish might resist ERVs-GTE integration because only 18 of the 367 species contained ERVs-GTE at relatively low copy numbers.

**Fig 2.**
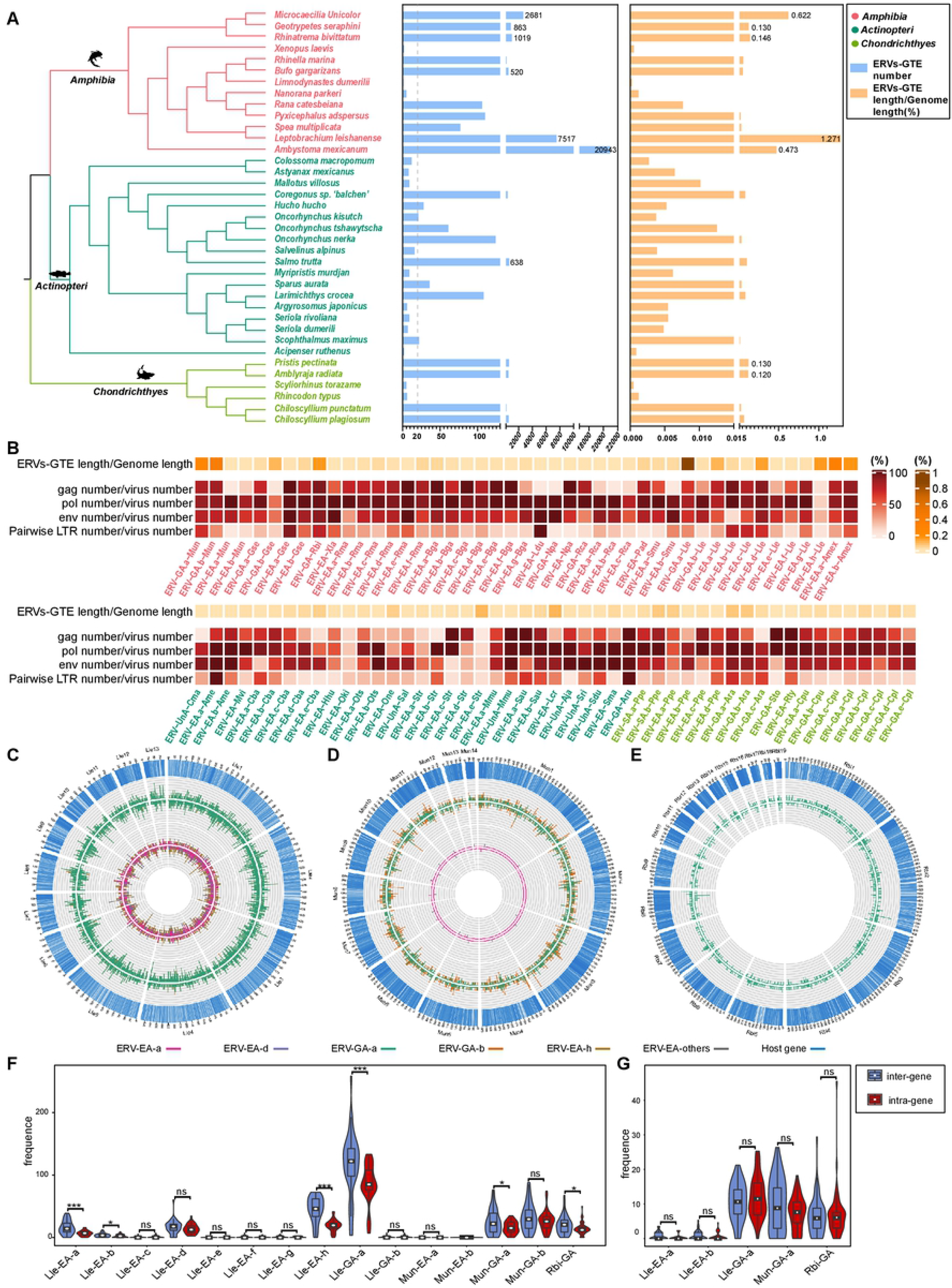
Distribution of ERVs-GTE in host genomes. (A) Copy numbers and proportions of ERVs-GTE in host genomes. The host phylogeny is obtained from TimeTree (http://www.timetree.org/) and shown on the left, and their classification is labeled using different colors. The copy numbers of ERVs-GTE higher than 500 are indicated accurately. ERVs-GTE lengths/genome lengths higher than 0.1% are indicated accurately. (B) Heatmap showing the proportion of major genes in each lineage. The host information of each viral lineage is labeled using different colors. The distribution of viral insertion in chromosomes of (C) *Leptobrachium leishanense,* (D) *Microcaecilia unicolor* and (E) *Rhinatrema bivittatum*. From inside to out, the concentric circles denote chromosome ideograms with gene regions represented as blue, followed by the ERV-GA intragene insertion frequency, ERV-GA intergene insertion frequency, ERV-EA intragene insertion frequency and ERV-EA intergene insertion frequency. The frequency was calculated as the viral insertion count within 1 Mb genomic ranges. The colors match the types of viral lineages. ERV-EA.b/c/e/f/g-Lle are grouped into ‘others’ due to their low frequency. Violin plot showing the distribution of (F) the viral insertion and (G) the young viral insertion (LTR identity > 99%) relative frequency by viral lineage. The relative frequency was calculated within a 100 Mb genomic range and then standardized against the total length of genes or non-genes within each species. Violin plot features: White point, median; box, 25th to 75th percentile; whiskers, data within 1.5× the interquartile range. Colors represent the location of insertions (intergene or intragene). The statistical analysis was carried out using the wilcoxon test. The data were considered statistically significant when *p ≤ 0.05, **p ≤ 0.01, and ***p ≤ 0.001.

Because some ERVs-GTE were too divergent to have been derived from the same viral lineage, we divided them into different lineages according to their genomic similarity. Accordingly, we found that 9 amphibians, 6 ray-finned fish and 4 cartilaginous fish harbored at least two lineages, and the number of lineages in different genomes varied from 1 to 8 (Fig 2B). Previous research indicated that ERVs without Env can more easily proliferate than those with Env [21]. However, ERVs-GTE with high copy numbers (>1000) tended to carry Env (66%-91%) rather than losing them, with the exception of ERV-EA.h-Lle, for which only 20% of copies contained Env. In addition, we also found that compared with major genes, pairwise long terminal repeats (LTRs) were harbored by relatively few ERVs-GTE. Notably, the copy number of ERV-EA.a- Amex was 14,680, and ERV-GA.a-Lle constituted 0.86% of the *L. leishanense* genome. To our knowledge, this finding constitutes the first time that the copy number of an ERV lineage was found to reach 10,000 and that an ERV lineage was found to constitute nearly 1% of its host genome (Fig 2B).

We found that ERVs-GTE in *A. mexicanum, L. leishanense, M. unicolor and R. bivittatum* constituted more than 0.1% of the host genomes, which was markedly higher than the proportion in other species (Fig 2B). To understand their distribution in host genomes, we generated an insertion map of ERVs-GTE in each genome (Fig 2C-E) and found that ERVs-GTE were widely distributed on all chromosomes and that some of them resided within genes or flanking genes. We then compared the intergene and intragene relative frequencies of viral insertions (Fig 2F). Seven lineages of ERVs-GTE, including 5 lineages in *L. leishanense*, 1 lineage in *M. unicolor* and 1 lineage in *R. bivittatum*, showed a strong tendency for intergene insertion rather than intragene insertion. However, their younger viral copies of these lineages (pairwise LTR divergence < 1%) were more likely to be randomly inserted (Fig 2G). Taking these findings into consideration, it appeared that the intragene insertion tendency of ERVs- GTE was the result of strong selection by the host rather than the virus’s own insertion preference.

### Characterization and classification of novel GTE

To better understand the relationship among all GTEs, an Env protein phylogenetic tree was generated using the reference GTEs and a representative GTE, which is the consensus or the longest and most completed GTE, in each lineage (Fig 3A, S1 Fig and S2 Data). The phylogeny revealed the diverse evolutionary status of GTEs because they formed at least 9 major clades, including 2 well-defined GTE-C.5-AV and GTE-C.9- GV clades. Most (90.1%) of our newly identified GTEs were in the well-supported Env-C.1 clade, which was distantly related to the well-defined AV and GV clades, whereas other identified GTEs were divided into 6 novel monophyletic groups, indicating their different evolutionary statuses. Additionally, gamma-type percomORFs previously identified in ray-finned fish [17] were clustered in Env-C.2 with newly identified GTEs in cartilaginous fish.

**Fig 3.**
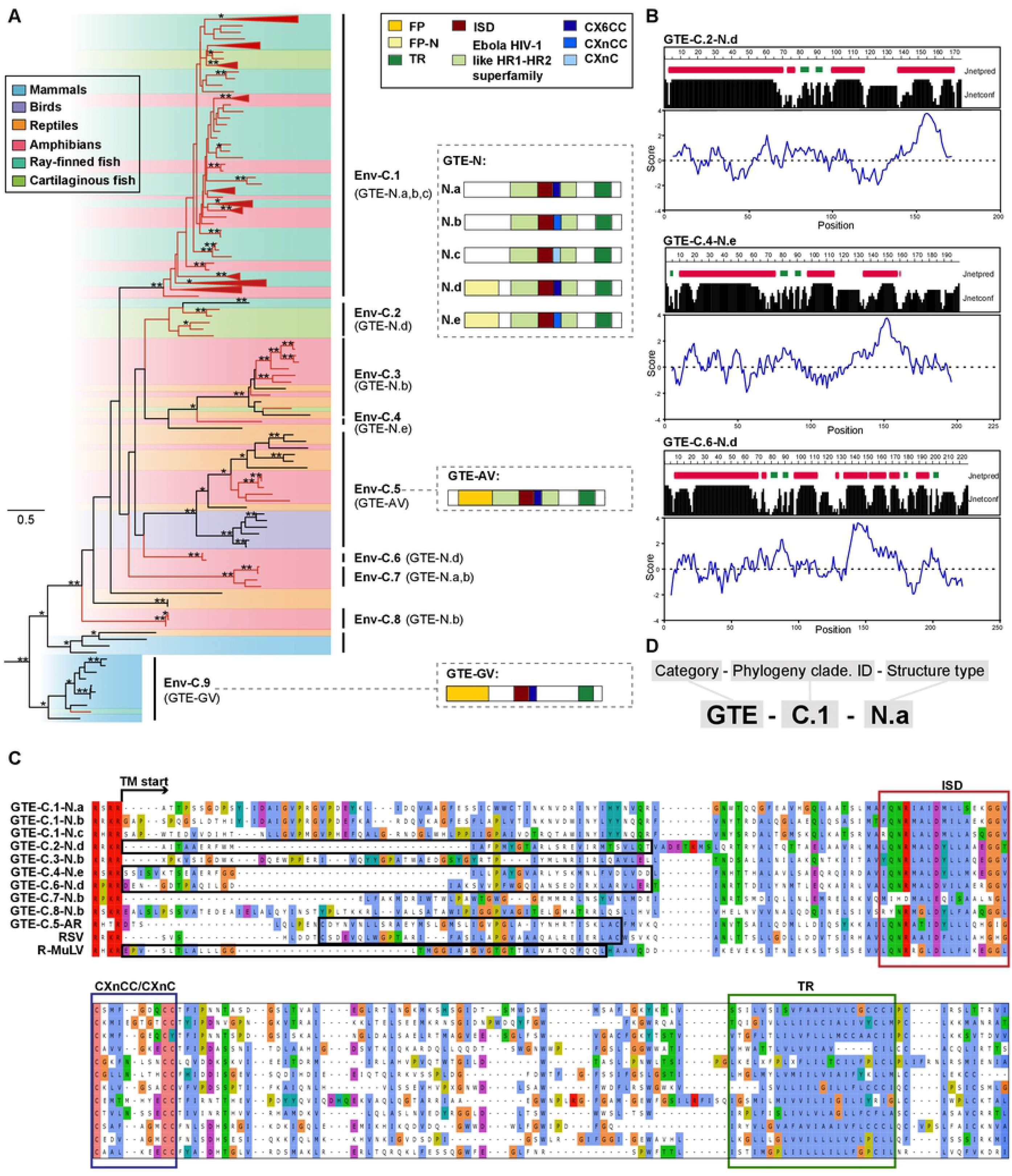
Characterization and classification of novel GTEs. (A) Phylogeny of GTEs. The left tree was inferred using amino acid sequences of the conserved region of transmembrane subunit (TM). The tree is rooted in deltaretroviruses, which are not shown in the tree. The newly identified ERVs-GTE and exRVs-GTE are labeled in red. The host information of each retrovirus is indicated using a shaded box. Bootstrap values lower than 65% are not shown. Single asterisks indicate values higher than 65%, while double asterisks indicate values higher than 80%. The scale bar indicates the number of amino acid changes per site. The typical GTE structures are shown on the right. The letters in the bracket besides the clade name indicate which GTE is found. FP, fusion peptide; ISD, immunosuppressive domain; TR, transmembrane region. CX6CC represents the three-cysteine motif of GTEs. (B) Prediction results of secondary structure and hydropathicity scores of amino acids in TM. Prediction results of GTE- C.2-N.d, GTE-C.4-N.e, GTE-C.6-N.d are shown here. The Secondary structure is shown at the top and the corresponding amnio acid hydrophilic index is shown at the bottom. Jnetpred is secondary structure prediction for TM. The alpha helix is labeled with red box and beta fold is labeled as green box. Jnetconf shows the reliability of prediction accuracy, range from 0 to 9, and large means more robust. The blue curve shows the calculated hydrophilic scores of amino acids, A posi- tive number means the protein is hydrophobic and vice versa. (C) Alignment of FP, ISD, the CX6CC motif and the flanking conserved domain of representative ERVs-GTE and other exogenous viruses. FP is labeled with black boxes. (D) The nomenclature for GTEs.

However, the phylogeny did not show a strong classification protocol for GTEs. Therefore, we then compared the distribution of motifs and domains in different GTEs.

Five major domains/motifs were found in GTEs: fusion peptide (FP), ISD, CX6CC, heptad repeat and transmembrane region (TR). By checking the structure of all GTEs, we found that ISDs and TRs were present in all GTEs, and heptad repeats could be identified in all GTEs except GTE-C.9-GV (Fig 3A). CX6CC had 3 subtypes, including CX6CC, CXnCC and CXnC, which were classified according to the functional residue distribution. The first two cysteines in the CX6CC motif participate in the formation of an intramolecular loop in the TM ectodomain, whereas the third cysteine is needed for formation of a covalent bond between the TM and SU domains [22]. Most (79.6%) GTEs contained CX6CC, whereas others contained CXnCC and CXnC. However, the newly identified GTEs showed no similarity to GTE-C.5-AV and GTE-C.9-GV in the N-terminus of the TM, where the FP is usually located. Because the FP mostly contains an alpha helical or beta structure and is hydrophobic [23], the secondary structure and hydrophobicity of the TM were used to predict the presence of potential novel FPs. This analysis led to the identification of 3 novel FPs that showed no sequence similarity to other viral FPs (Fig 3C). By taking all of these findings into consideration, the structure of GTEs can be mainly classified into 7 types, as shown in Figure 3A. To better illustrate the difference among all the GTEs, alignments of all major groups of GTEs are shown, which reconfirmed the conservation of the ISD, CX6CC and TR and the diversity of FPs (Fig 3C). GTEs cannot be classified by phylogeny alone. Here, by combining the phylogeny and TM structure, we proposed a nomenclature for GTEs (Fig 3D).

### Extensive recombination of ERVs-GTE

To further elucidate the relationship among ERVs-GTE, exRVs-GTE and those from other RVs, a phylogenetic analysis based on Pol (>600 aa) was performed (S3 Fig and S3 Data). This phylogeny revealed that our viral elements could be divided into three different groups: 1) Class I epsilon-related viruses, which can be divided into three clades, namely, Pol-LE.1, Pol-LE.2 and epsilon; 2) Class I gamma-related viruses, which can also be divided into three clades, namely, Pol-LG.1, Pol-LGE.1 and Gamma; and 3) Class III SnRV-related viruses. The pol and env phylogenies were then compared, and the results indicated that recombination was widespread and frequently occurred among different RVs over long evolutionary timescales.

The genomic structure also reflected recombination among viruses. Epsilonretroviruses are typical complex RVs that encode accessory genes, and their Env proteins are not GTEs or BTEs [13]. However, our epsilon-related class I ERVs contained GTEs instead of epsilon Envs and did not carry any accessory gene, indicating potential recombination between epsilonretrovirus and ancient RVs-GTE (Fig 4B and S4 Fig). In contrast, our gamma-related class I ERVs were much more highly conserved because they exhibited a typical GV structure and contained all the conserved domains/motifs carried by exogenous RVs (e.g., R-MuLV).

**Fig 4.**
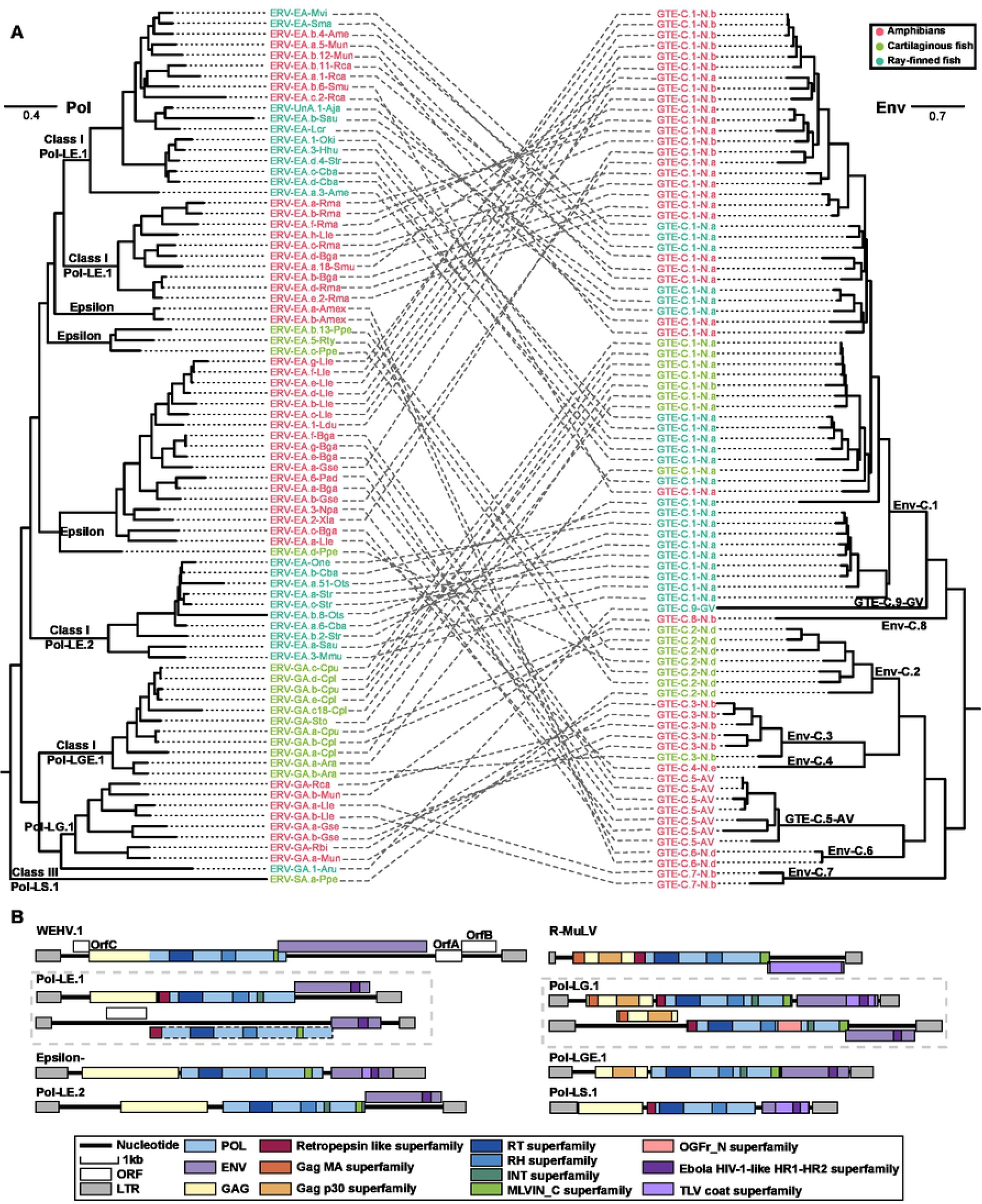
Genetic exchange of ERVs-GTE. (A) Phylogenetic incongruence between Pol and Env of ERVs-GTE. The classification of Pol and Env is shown at each node. (B) The comparison of genomic structure of exogenous retroviruses and representative ERVs-GTE. The WEHV (epsilonretrovirus) and R-MuLV (gammaretrovirus) genomes are shown on the top. The predicted domain or region that encodes conserved proteins is labeled with colored boxes. The dotted boxes indicate putative open reading frames (ORFs) containing stop codons or indels.

### Conserved domain shared between GTEs and other divergent viruses

Filoviruses (ssRNA(-)), particularly Ebola virus, were previously discovered to share a similar ISD and CX6CC motif with GTEs [24]. To search for other possible gene sharing events among different viruses, we used all the discovered novel GTEs to perform a search against the NCBI Nr database and surprisingly found that F-env on HP35 of chelonid alphaherpesvirus 5 (a DNA virus) [25] shared 26.68% similarity and 92% coverage with ERV-GA.a-Ara and 31.75% similarity and 68% coverage with ERV-GA.a-Cpu. By searching against the conserved domain database (CDD), we found that this protein contained two conserved domains: 1) the Ebola HIV-1-like HR1-HR2 superfamily domain (cl02885), a typical retroviral domain that plays a key role in the dynamic rearrangement of the trimer during the process of fusion [11, 17], and 2) the TLV coat superfamily domain (cl27694) (Fig 5A). An alignment comparison indicated that chelonid alphaherpesvirus 5 also contained homologous ISD, CX6CC motif, TR and flanking conserved regions with lengths of 164 aa (Fig 5B and S6 Data) but showed no significant similarity in other regions, including the FP. However, other herpesviruses were also screened, and we found that they did not contain such domains and motifs. This finding indicated that the transfer of ISD, CX6CC and TR could be a one-off event in reptiles.

**Fig 5.**
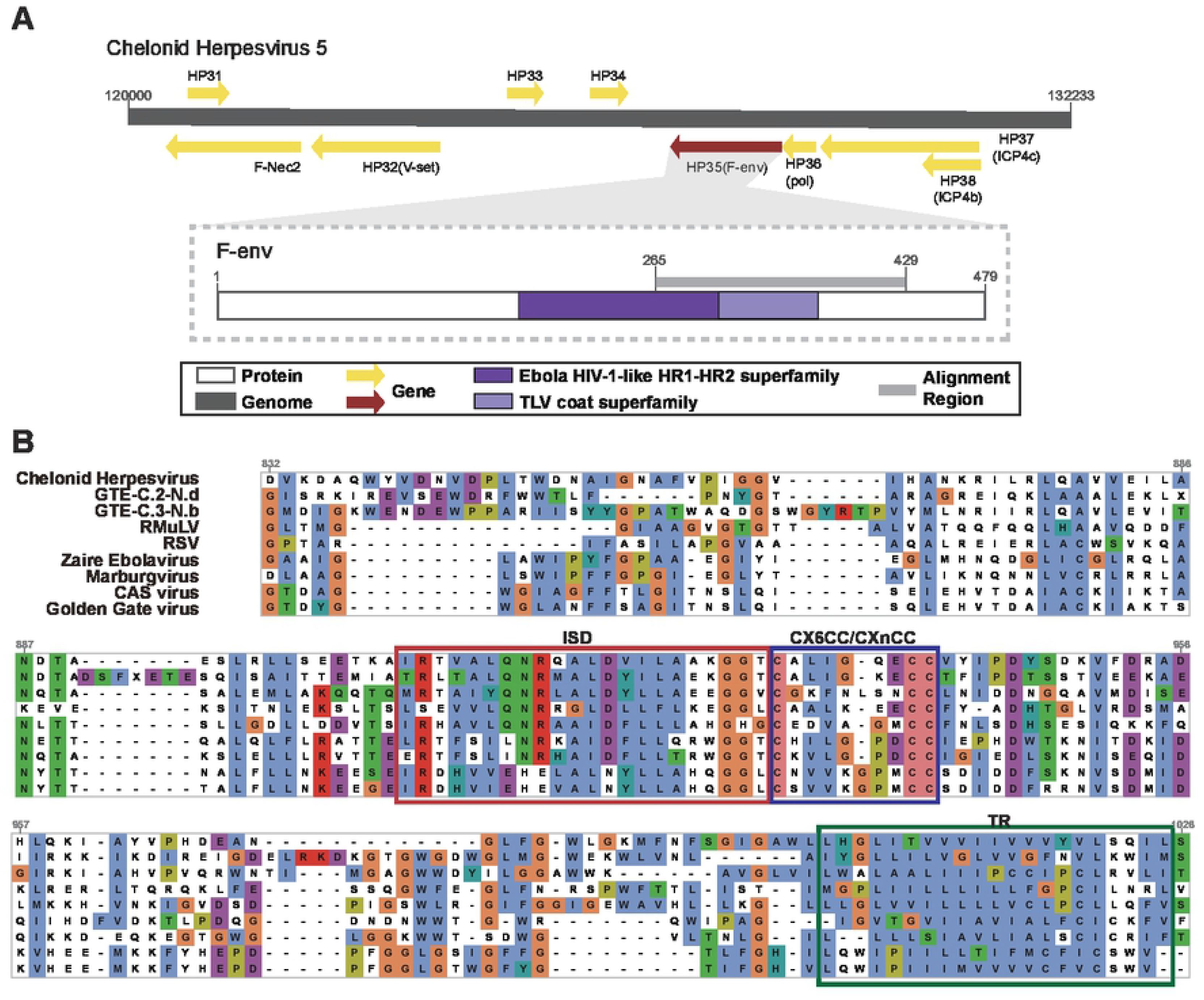
Conserved domain sharing among divergent viruses. (A) Partial genomic map of Chelonid Herpesvirus and genomic structure of F-env in Chelonid Herpesvirus. The predicted open reading frames (ORFs) or region that encode conserved proteins is labeled with colored arrows or boxes. (B) Alignment of FP, ISD, the CX6CC motif and the flanking conserved domain of Chelonid Herpesvirus, ERVs-GTE which are most homolog to Chelonid Herpesvirus, and other exogenous viruses.

### The host gene was captured by RVs-GTE in amphibians

The opioid growth factor receptor (OGFr) gene, which plays an important role in the regulation of cell growth and embryonic development, is a typical animal gene [26]. Unexpectedly, by using CD-Search, we found that 51.2% ERV-GA.a-Lle, 41.2% ERV- GA.b-Lle, 23.1% exRVssi-GTE and 25% exRVlbo-GTE were all found in *Megophryidae*, and all of these encoded OGFr within their Pol gene ORF (Fig 6A). By checking the structure of viral OGFr, we found that they only contained exons. To further elucidate the relationship between host OGFr and viral OGFr, a phylogenetic tree of all related amphibians and viral OGFr was constructed (Fig 6B). This tree revealed that viral OGFr formed a monophyletic clade, which was distant from the OGFr of their hosts.

**Fig 6.**
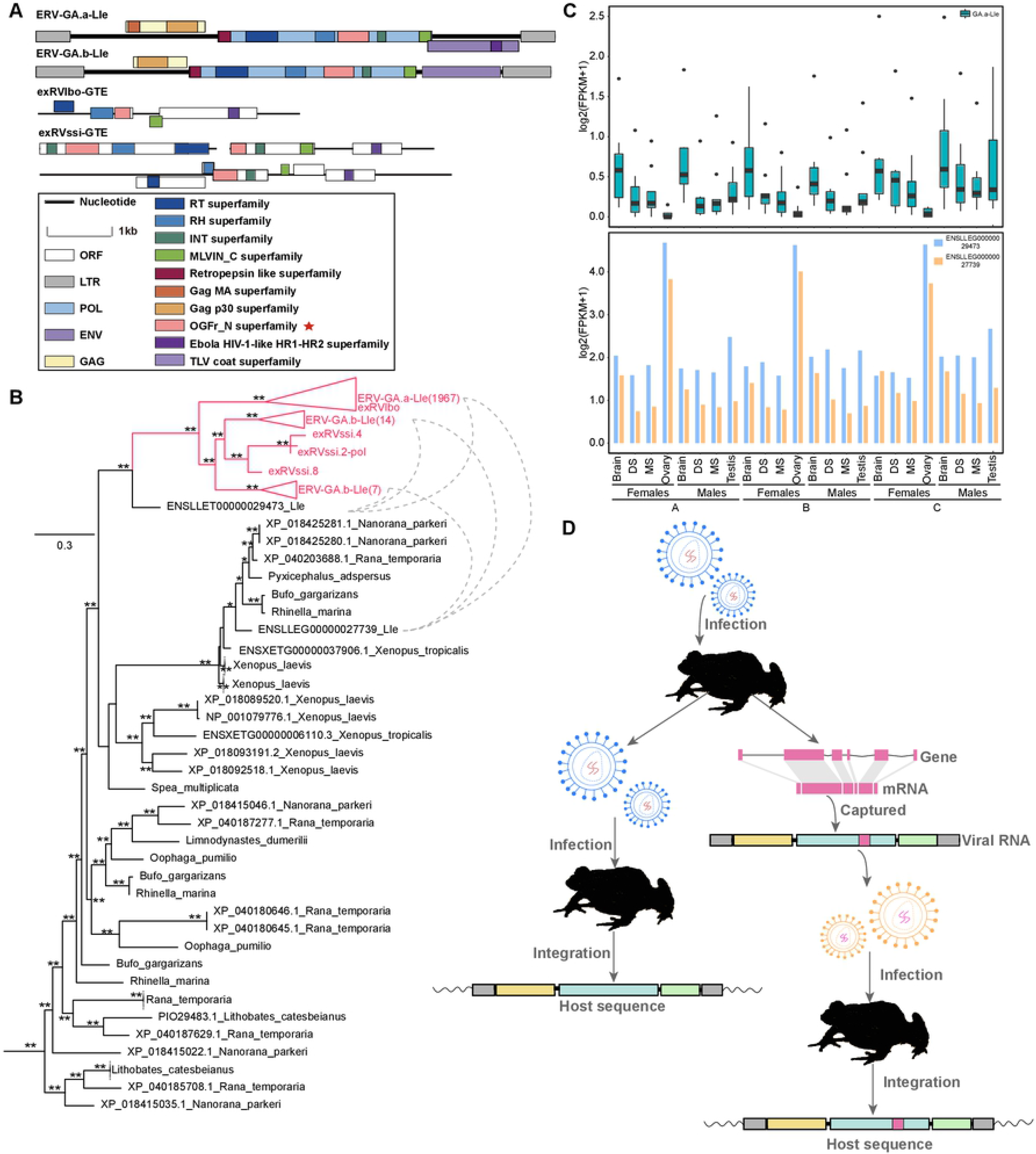
Host gene capturing by RVs-GTE. (A) Genomic structure of ERVs-GTE and exRVs-GTE contained opioid growth factor receptor gene (OGFr). The predicted domain or region that encodes conserved proteins is labeled with colored boxes. Red asterisks indicate that the motif does not have a viral origin. (B) Phylogeny of viral and frog OGFr domain. Viral OGFr were labeled in red and were linked by dotted lines with OGFr of their animal hosts. (C) The expression level of viral and host OGFr in *L. leishanense.* The expression level was evaluated by log_2_(FPKM+1). Box plot of the expression of viral OGFr was shown at the top, while the bar chart for the expression of host OGFr was shown at the bottom. For each boxplot of viral OGFr, the loci expression values were calculated based on three biological replicates (S7 Data). The middle black box indicates the median of the expression values. The upper and lower terminal lines of a box represent the 25th and 75th percentiles, respectively. Lines extending vertically from the boxes (whiskers) indicate variability outside the upper and lower percentiles. All the sequenced transcriptomes from *L. leishanense* were divided into 2 genders and 3 developmental stages. A: the subadult period, B: the adult breeding period, C: the postbreeding stage. DS: dorsal skin; MS: maxillary skin. (D) Evolution scenario of retrovirus capturing host OGFr.

Furthermore, we also found that some ERVs-GA-Lle contained intact ORFs, which indicated that they had the potential to express. Accordingly, we collected all 72 available transcriptome data of *L. leishanense* to further investigate their expression and compared them with the host OGFrs. We found that the minority (9/2313) of the OGFrs of ERV-GA.a-Lle were expressed, whereas all OGFrs of ERV-GA.b-Lle were barely expressed. Additionally, 2 OGFrs of ERV-GA.a-Lle located on chromosome 6 were expressed at a markedly higher level than the others (S7 Data). Overall, both viral and host OGFrs were expressed in different tissues, and the expression of the latter was higher than that of the former across all tissues (Fig 6C). Interestingly, the host OGFr was highly expressed in the ovary, whereas the viral OGFr was seldom expressed, indicating their potential interaction in such tissue.

By taking these findings into consideration, OGFr capture more likely occurred during infection with the ancient exogenous ancestor RVs-GTE of these viruses occasionally than during infection with these viruses independently (Fig 6D).

### Estimation of the time of ERVs-GTE insertion

We also used vertical transmission to infer the ERVs-GTE insertion time. By comparing the flanking sequences and LTR similarity, we found 46 and 2 orthologous ERV groups in the genera *Chiloscyllium* and *Seriola*, respectively (S3 Table). However, no divergence time could be found for the genus *Chiloscyllium*. We were only able to estimate the insertion of the latter group, which occurred 4.4∼22.2 MYA (S6 Fig).

## Discussion

In this study, we developed a phylogenomic approach by regarding env genes as seeds for the detection of ERVs and exRVs. Instead of using pol alone, we traced the evolutionary histories of RVs by combining pol and env, which allowed us to observe many fine distinctions and unique patterns during the macroevolution of RVs. We found that GTEs were widely distributed in vertebrates, and many of these showed diversity in well-defined GTE-C.5-AV and GTE-C.9-GV, as reflected by their phylogeny and genomic motif/domain distribution (Fig 3). On the one hand, the majority of GTEs of anamniotic RVs were found to lie on Env-C.1, which is phylogenetically divergent from the GTEs of amniotic RVs, showing host restriction regarding the transmission of viral progenitors of RVs-GTE. On the other hand, GTEs can be placed into different phylogenetic clades, representing the dynamics of the evolutionary status of GTEs. RVs in different classes of hosts could also share similar GTEs, showing the occasional possibility of cross-class transmission. Furthermore, numerous incongruences were observed in the Env and Pol phylogenies, which implied the occurrence of frequent recombination involving various RVs (Fig 4A). For example, ERV-SA.a-Ppe harbored Pol of Class III and GTE of Class I, as has also been observed previously [27]. By taking these aspects into consideration, our studies suggest that the Env analyses provide additional information that cannot be obtained via Pol analyses and thus expanded our understanding of the complex macroevolution of RVs.

Our studies also suggest the genomic complexity and elasticity of GTEs. First, the motifs/domains in GTEs tend to be highly variable. By combining the second structure and hydrophobicity predictions, 3 novel predicted retroviral FPs were identified in this study, and these showed no similarity to any other viral FP, as determined by BLASTp (Fig 3). This finding could reflect the possible multiple origins of GTE segments and indicate the limitation of a similarity-based search for viral counterpart identification. Addit ionally, three major variants of CX6CC have also been characterized based on consensus viral GTE comparisons and the function of each cysteine. However, we admit that further experimental confirmation is needed for such identification. Second, the motif/domain distribution of GTEs is more flexible. GTE mainly has 5 major motifs/domains, including the FP, heptad repeat, ISD, CX6CC and TR. In this study, we found that the ISD, CX6CC and TR are relatively conserved and carried by nearly all GTEs, whereas the FP and intact heptad repeats are not necessary counterparts. However, these domains play crucial roles during the process of fusion [28–32]. Overall, these findings indicate that GTEs may have their own complex evolutionary history, which involves multiple internal recombination events of different regions and results in the observed diversity.

Both GTEs and BTEs were screened in this study. However, no BTE was found, which may be due to its narrow host range (mammals) [11], whereas a large amount of GTEs were found in amphibians and fish in an unbalanced manner (S1 Data and Fig 2). Most importantly, the majority of amphibians harbored ERVs-GTE, and their copy numbers were higher than those in other vertebrates (Fig 2). Because previous research also found that many amphibians harbor other rare ERVs [20, 33], amphibians appear to be better reservoirs for RVs than other classes of vertebrates. However, further study is merited because only 19 amphibian genomes and 61 transcriptomes are currently available.

ERVs are usually present in the host genome at numbers ranging from a few to a few thousand [7, 34]. Surprisingly, we found that the copy number of ERV-EA.a-Amex reached nearly 15,000 and that ERV-GA.a-Lle constituted 0.86% of the host genome; both of these ERVs were markedly more abundant than all the ERVs discovered previously [7, 34–36]. Contrary to previous research, the majority of their copies (14680 for Amex, 4552 for a-Lle) contained env, which suggested that these ERVs may likely proliferate by reinfection mediated by the Env rather than retrotransposition. Because ERVs-GTE with high copies were recombinant viral elements carrying different types of pol and the same GTE, it appears that recombinant RVs, particularly those carrying GTEs, were key to infection and were well adapted to their hosts.

Horizontal gene transfer from RVs to hosts is widely distributed and well documented. However, this transfer is rare between unrelated viruses with distinct genome types. Homologous GTEs have been found in both arenaviruses and filoviruses [14, 24, 37], and herpesviruses can also acquire RV genes (superantigens (sags)) [38]. Here, we found that chelonid alphaherpesvirus 5 can also harbor a similar partial GTE that contains intact ISD and CX6CC (Fig 5 and S6 Data). However, no other herpesviruses were found to contain such domains or motifs. Herpesviruses typically infect hosts indefinitely and move from latent to lytic replication during periods of immunosuppression [38]. A host infected with RVs is more likely to be in an immunocompromised state that provides an ideal environment for gene transfer in an activated and replicating herpesvirus. Because reptiles can be infected by both RVs- GTE and herpesviruses, the acquisition of ISD and CX6CC motifs could be an occasional horizontal gene transfer event that occurs during coinfection in reptiles. In addition, the results further reconfirm that gene exchange can occur among divergent viruses [38, 39].

Numerous experimental studies have confirmed that RVs can be coopted by their host [40–43]. Here, we reconfirmed that host genes could also be captured by RVs-GTE [44] and fully depicted the evolutionary scenario of such events (Fig 6D). Four viral lineages in 3 hosts were confirmed to carry OGFrs. In regard to ERV-GA.a-Lle and ERV-GA.b, 2331/4552 and 21/51 of the copies contained the intronless OGFr gene, which indicates that gene capture occasionally occurred in some RVs-GTE during infection rather than their endogenization. We also found that most of these had intact env genes, indicating that the proliferation of such viruses was more likely to be caused by repeated infection rather than transposons. OGFr is thought to be related to cancer because previous studies have indicated that the upregulation of OGFr represses the growth of cancer cells in culture and in nude mice [45]. On the one hand, the proliferation of OGFr could help the host repress cancer and could prolong the lifespan of the host. On the other hand, OGFr might help RVs produce and proliferate within hosts. It appears that the ‘co- option’ of OGFr is a win-win situation for both hosts and viruses. Such events benefit our understanding of the interaction between the host and viruses. However, we also admit that further studies on the functional influence of viral OGFr on animal hosts is warranted.

The transcriptome data provided us with additional resources for discovering novel exogenous and endogenous viruses. Regarding their flaws, on the one hand, major genes (gag, pol, and env) of exRVs-GTE reside in different contigs and cannot be assembled at the genomic level in most cases. Therefore, it is rather difficult to characterize recombination among different viral elements. On the other hand, the efficacy of viral discovery through transcriptomic analysis is highly dependent on the sequencing depth and assembly quality. Thus, the absence of viral contigs in specific species cannot be explained by the noninfectious nature of this virus. Regardless of this fact, the pipeline presented in this manuscript provides a refined procedure for discovering novel exRVs.

In summary, by integrating genomics and transcriptomic data and incorporating a multiple-round screening method and phylogenomic analysis, we revealed the hidden diversity and genomic elasticity of GTEs, which were found to be widely distributed in different vertebrates. Additionally, recombinant ERVs-GTE in several amphibians proliferated rampantly and under strong host selection. The incongruence between Env and Pol phylogenies suggested that recombination frequently occurred between different RVs across taxonomic barriers. These findings demonstrate the feasibility and practicability of using Env to perform a phylogenetic analysis and reveal hidden evolutionary features, which indicate the complex macroevolutionary history of RVs.

## Materials and Methods

### Genome and transcriptome screening and identification of ERVs-GTE and exRVs-GTE

To discover potential targeted viral elements in fish and amphibian genomes, all 974 available fish and 19 amphibian genomes (S1 Table) were first screened using the tBLASTn algorithm [46], and the Env proteins of all reference RVs-GTE and their endogenous forms (S2 Table) were used as probes. A 40% sequence identity over 40% of the region with an e-value of 1E-5 was used to filter significant hits. Second, the significant hits confirmed by phylogeny were concatenated based on the host genomic location and alignment positions with reference RV-GTE proteins. Third, the same genomes were subjected to a second round of screening by tBLASTn using the protein sequences of concatenated Env. Finally, the flanking sequences of significant hits obtained from the second round that were confirmed by phylogeny were extended to identify viral pairwise LTRs using BLASTN [46], LTR_Finder [47] and LTR_harvest [48]. Full-length ERVs were used as queries to search for ERV copies using BLASTn. The hit parts of sequences longer than 3 kb with 80% identity were regarded as copies of each ERVs-GTE lineage (S1 Data), and these were named in accordance with the previous nomenclature proposed for ERVs [8].

To identify potential exRVs, all 439 datasets in the transcriptome sequencing assembly database (TSA) (S1 Table) were screened using tBLASTn, and the Env proteins of the reference RVs-GTE, including the newly identified ERVs-GTE, were used as probes. A 40% sequence identity over 40% of the region with an e-value of 1E-5 was used to filter significant hits. The called hit contigs were then included in the phylogenetic analysis. The viral contigs within the GTE clade were considered.

### Consensus genome construction and genome annotation

ERVs-GTE longer than 5 kb in each lineage were aligned using MAFFT 7.222 [49] and then used to construct consensus sequences for each EGV lineage by Geneious. The distributions of ORFs in copies of EGV and exGV contigs were determined using ORFfinder (https://www.ncbi.nlm.nih.gov/orffinder/) in the NCBI database and confirmed by BLASTp [46]. Conserved domains for each sequence were found using CD-Search against the CDD (https://www.ncbi.nlm.nih.gov/cdd/) [50]. The consensus sequences of ERVs-GTE can be found in S4 Data. The TM of the Env protein of representative ERVs-GTE in each type was used to predict the secondary structure and hydropathicity scores with JPred 4 (http://www.compbio.dundee.ac.uk/jpred/) and ExPASy (https://web.expasy.org/protscale/). The predicted results are shown in Fig 3B and S2 Fig.

### Construction of the ERVs-GTE insertion site map

The distribution of the viral integration, which was classified as intragene or intergene, was defined depending on whether an overlap existed between an inserted virus and a gene extended by 10 kb on both sides. BEDtools [51] was used for overlap identification and to determine the frequency per 1000 kb of the genome. Circos [52] was used for visualization of the results.

### Phylogenetic analysis

To investigate the evolutionary relationship between the RVs and viral elements found in this study, the protein sequences of the TM region, Env and Pol were aligned using MAFFT 7.222 [49]. The regions in the alignment that aligned poorly were removed using TrimAL [53] and confirmed manually with MEGA X [54]. A sequence was excluded if its length was less than 75% of the alignments. The best-fit models (Env: LG+G4; Pol: LG+F+I+G4) were selected using ProtTest [55], and phylogenetic trees for these protein sequences were inferred using the maximum likelihood (ML) method in IQ-Tree (Env and Pol phylogeny) [56] by incorporating 100 bootstrap replicates for the assessment of node robustness. The phylogenetic trees were viewed and annotated using FigTree V1.4.3 (https://github.com/rambaut/figtree/). The alignments performed in this study can be found in S2 Data and S3 Data.

### Examination of the recombination of ERVs-GTE

The Pol and Env sequences of representative ERVs-GTE were used to construct corresponding ML phylogenetic trees. The regions in the alignment that aligned poorly were removed using TrimAL [53] and confirmed manually with MEGA X [54]. The best-fit models (Env: LG+I+G4; Pol: LG+F+G4) were selected using ProtTest [55], and the phylogenetic trees for these protein sequences were inferred using the maximum likelihood (ML) method in IQ-Tree [56] by incorporating 100 bootstrap replicates for the assessment of node robustness. The phylogenetic trees were viewed and annotated using FigTree V1.4.3 (https://github.com/rambaut/figtree/). The phylogenetic tree match between Pol and Env was estimated manually.

### Dating analysis for determining the integration time of ERVs-GTE

To determine the potential vertical transmission among all ERVs-GTE, we first extracted the flanking sequences of ERVs-GTE with lengths of 1 kb at both sites. The sequences on both sides of an ERVs-GTE and 300 bp of its LTRs were then concatenated and compared with each other using dc-megablast with an e-value of 1E-5. Significant pairwise sequences were retrieved if the alignment coverage was over 50% and longer than 800 bp.

### Gene expression analysis

Seventy-two *L. leishanense* RNA-seq samples [57], which are available in the Sequence Read Archive (SRA) database (SRR8736149–SRR8736220), were aligned to the reference genome (ASM966780v1) using HISAT2 [58]. The expression levels were then computed for each group of samples after filtering the loci with low expression (more than 20 samples had no expression). The OGFr expression level in each RNA-seq library was calculated as fragments per kilobase million (FPKM).

## Data availability

All the data needed to support the conclusions detailed in the article are included in the article itself and the supplementary data.

## Acknowledgement

This work was supported by National Natural Science Foundation of China (31970176) and CAS Pioneer Hundred Talents Program to J.C.

## Author contribution

Y.C. and J.C. designed research; Y.C., X.W., M.L. and Y.S. performed research; Y.C., X.W., M.L. and Y.S. analyzed data; and Y.C., X.W. and J.C. wrote the paper.

## Supporting information

**S1 Fig. Phylogenetic analyses and characterization of envelope structure of ERVs- GTE, exRVs-GTE and exogenous retroviruses.** (Left) The envelope phylogeny tree of exogenous retroviruses, ERVs-GTE and exRVs-GTE. The trees were inferred using amino acid sequences of transmembrane domain (TM) and flanking conserved regions of Env gene. Bootstrap values lower than 65% are not shown. Single asterisks indicate values higher than 65%, while double asterisk indicate values higher than 80%. The scale bar indicates the number of amino acid changes per site. (Right) Distribution of CXXC motif, FP (fusion peptide), Alpha_FP (fusion peptide flanked by a pair of cysteines), ISD (immunosuppressive domain) and CX6CC motif in envelopes of exogenous retroviruses, ERVs-GTE and exRVs-GTE.

**S2 Fig. Prediction of secondary structure of TM in ERVs-GTE.** The predicted secondary structure and hydrophobicity of transmembrane domain (TM) in GTE-C.1-N.a (A), GTE-C.3-N.b (B), GTE-C.7-N.b (C), GTE-C.8-N.b (D), GTE-C.5-AR (E), GTE-C.9-GR (F) and ZFERV (G) are shown. The Secondary structure is shown at the top and the corresponding amnio acid hydrophilic score is shown at the bottom. Jnetpred is the secondary structure prediction for TM. The alpha helix is labeled with red box, while the beta fold is labeled with green box. Jnetconf shows the reliability of prediction accuracy, range from 0 to 9, and large means more robust. The blue curve shows the calculated hydrophilic scores of amino acids, A positive number means the protein is hydrophobic and vice versa.

**S3 Fig. Characterization of the conservation of ERVs-GTE and exRVs-GTE.** The pol phylogenetic tree of exogenous retroviruses, ERVs-GTE and exRVs-GTE. The trees were inferred using the amino acid sequences of the pol gene. The tree is rooted in foamy viruses, which are not shown in the tree. The newly identified viral elements are labeled in bold. The host information of each retrovirus is indicated using a shaded box. Bootstrap values lower than 65% are not shown. Single asterisks indicate values higher than 65%, while double asterisks indicate values higher than 80%. The scale bar indicates the number of amino acid changes per site.

**S4 Fig. Representative genomic structures of ERVs-GTE and exRVs-GTE.** Representative genomic structure of ERVs-GTE (A) and exRVs-GTE (B). Different classes are marked with colored fonts. The predicted domain or region that encode conserved proteins are labeled with colored boxes. The dotted boxes indicate putative open reading frame (ORF) for containing stop codons or indels. Red asterisk indicates the motif which does not have a viral origin.

**S5 Fig. OGFr_N super family domain alignments of the ERVs-GTE and exRVs- GTE in *Megophryidae.*** The conserved domains were determined by searching the Conserved Domain Database (CDD). (A) The alignments of ERVs-GTE in *Leptobrachium leishanense* genome and OGFr_N super family (pfam04664); (B) The alignments of exRVs-GTE in the transcriptome data of *L. boringii*, *Scutiger cf. sikimmensis* and OGFr_N super family (pfam04664). Numbers refer to the position in the ERVs-GTE protein (exRVs-GTE nucleotides) or conserved domain. Identical amino acid residues are highlighted in red, and grey and blue indicate gaps or different amino acid residues, respectively. The E-value was generated by Conserved Domain search.

**S6 Fig. ERVs-GTE orthologous insertions in *Chiloscyllium* and *Seriola*.** (A) ERVs- GTE integration events during fish evolution. The species tree was constructed by TimeTree. Different classes are marked with colored fonts. Arrowheads indicate the events of ERVs-GTE integrations. Integration events are labeled numerically. Dotted lines only indicate the genetic relationship and do not imply the divergence time of these species. (B) Examples of orthologous insertions for ERVs-GTE. Rectangles from left to right represent 1,000 bp flanking sequence, 5’-LTR, internal genes of an ERV, 3’- LTR, and 1,000 bp flanking sequence, respectively. Dashed boxes indicate missing of the corresponding regions.

**S7 Fig. Genetic distance of TM in representative ERVs-GTE and exogenous viruses.** The Heatmap shows the p-distance among representative ERVs-GTE and exogenous viruses in TM conserved region. Chelonid alphaherpesvirus-5 is labeled in orange, while newly identified ERVs-GTE are labeled in red.

**S1 Table. Information of fish and amphibian genomes and transcriptome sequence assembly (TSA) database used for data mining.**

**S2 Table. Information on the representative retroviruses.**

**S3 Table. Information of the ERVs-GTE orthologous insertions in *Chiloscyllium* and *Seriola*.**

**S1 Data. Information of all ERVs-GTE and exRV-GTE in this study.**

**S2 Data. Env alignment of ERVs-GTE, exRVs-GTE and exogenous ERVs.**

**S3 Data. Pol alignment of ERVs-GTE, exRVs-GTE and exogenous ERVs.**

**S4 Data. Consensus sequences of ERVs-GTE.**

**S5 Data. Alignment of representative ERVs-GTE and other retroviruses.**

**S6 Data. Alignment of representative ERVs-GTE and exogenous viruses.**

**S7 Data. Reads counts of OGFr in ERV-GA.a-Lle.**

## References

1. Rous P. A SARCOMA OF THE FOWL TRANSMISSIBLE BY AN AGENT SEPARABLE FROM THE TUMOR CELLS. J Exp Med. 1911;13(4):397–411.

2. Hahn BH, Shaw GM, De Cock KM, Sharp PM. AIDS as a zoonosis: scientific and public health implications. Science. 2000;287(5453):607–14.

3. Weiss RA. Retroviruses and human cancer. Semin Cancer Biol. 1992;3(5):321–8.

4. Xu W, Eiden MV. Koala Retroviruses: Evolution and Disease Dynamics. Annu Rev Virol. 2015;2(1):119–34.

5. Johnson WE. Origins and evolutionary consequences of ancient endogenous retroviruses. Nat Rev Microbiol. 2019;17(6):355–70.

6. Stoye JP. Studies of endogenous retroviruses reveal a continuing evolutionary saga. Nat Rev Microbiol. 2012;10(6):395–406.

7. Xu X, Zhao H, Gong Z, Han GZ. Endogenous retroviruses of non-avian/mammalian vertebrates illuminate diversity and deep history of retroviruses. PLoS Pathog. 2018;14(6):e1007072.

8. Gifford RJ, Blomberg J, Coffin JM, Fan H, Heidmann T, Mayer J, et al. Nomenclature for endogenous retrovirus (ERV) loci. Retrovirology. 2018;15(1):59.

9. Henzy JE, Gifford RJ, Johnson WE, Coffin JM. A novel recombinant retrovirus in the genomes of modern birds combines features of avian and mammalian retroviruses. J Virol. 2014;88(5):2398–405.

10. Chen M, Cui J. Discovery of endogenous retroviruses with mammalian envelopes in avian genomes uncovers long-term bird-mammal interaction. Virology. 2019;530:27–31.

11. Henzy JE, Johnson WE. Pushing the endogenous envelope. Philos Trans R Soc Lond B Biol Sci. 2013;368(1626):20120506.

12. Chen Y, Chen M, Duan X, Cui J. Ancient origin and complex evolution of porcine endogenous retroviruses. Biosafety and Health. 2020;2(3):142–51.

13. Henzy JE, Coffin JM. Betaretroviral envelope subunits are noncovalently associated and restricted to the mammalian class. Journal of virology. 2013;87(4):1937–46.

14. Bénit L, Dessen P, Heidmann T. Identification, phylogeny, and evolution of retroviral elements based on their envelope genes. Journal of virology. 2001;75(23):11709–19.

15. Chen M, Guo X, Zhang L. Unexpected discovery and expression of amphibian class II endogenous retroviruses. Journal of virology. 2021;95(3):e01806–20.

16. Niewiadomska AM, Gifford RJ. The extraordinary evolutionary history of the reticuloendotheliosis viruses. PLoS Biol. 2013;11(8):e1001642.

17. Henzy JE, Gifford RJ, Kenaley CP, Johnson WE. An Intact Retroviral Gene Conserved in Spiny-Rayed Fishes for over 100 My. Mol Biol Evol. 2017;34(3):634–9.

18. Shen CH, Steiner LA. Genome structure and thymic expression of an endogenous retrovirus in zebrafish. Journal of virology. 2004;78(2):899–911.

19. Shi J, Zhang H, Gong R, Xiao G. Characterization of the fusion core in zebrafish endogenous retroviral envelope protein. Biochemical and biophysical research communications. 2015;460(3):633–8.

20. Chen Y, Zhang YY, Wei X, Cui J. Multiple Infiltration and Cross-Species Transmission of Foamy Viruses across the Paleozoic to the Cenozoic Era. Journal of virology. 2021;95(14):e0048421.

21. Magiorkinis G, Gifford RJ, Katzourakis A, De Ranter J, Belshaw R. Env-less endogenous retroviruses are genomic superspreaders. Proc Natl Acad Sci U S A. 2012;109(19):7385–90.

22. Greenwood AD, Ishida Y, O’Brien SP, Roca AL, Eiden MV. Transmission, Evolution, and Endogenization: Lessons Learned from Recent Retroviral Invasions. Microbiol Mol Biol Rev. 2018;82(1).

23. Del Angel VD, Dupuis F, Mornon JP, Callebaut I. Viral fusion peptides and identification of membrane-interacting segments. Biochem Biophys Res Commun. 2002;293(4):1153–60.

24. Gallaher WR. Similar structural models of the transmembrane proteins of Ebola and avian sarcoma viruses. Cell. 1996;85(4):477–8.

25. Ackermann M, Koriabine M, Hartmann-Fritsch F, de Jong PJ, Lewis TD, Schetle N, et al. The genome of Chelonid herpesvirus 5 harbors atypical genes. PLoS One. 2012;7(10):e46623.

26. Guo Y, Wang L, Zhou Z, Wang M, Liu R, Wang L, et al. An opioid growth factor receptor (OGFR) for [Met5]- enkephalin in Chlamys farreri. Fish Shellfish Immunol. 2013;34(5):1228–35.

27. Yedavalli VRK, Patil A, Parrish J, Kozak CA. A novel class III endogenous retrovirus with a class I envelope gene in African frogs with an intact genome and developmentally regulated transcripts in Xenopus tropicalis. Retrovirology. 2021;18(1):20.

28. Epand RM. Fusion peptides and the mechanism of viral fusion. Biochimica et Biophysica Acta (BBA) - Biomembranes. 2003;1614(1):116–21.

29. Apellaniz B, Huarte N, Largo E, Nieva JL. The three lives of viral fusion peptides. Chem Phys Lipids. 2014;181:40–55.

30. Mzoughi O, Teixido M, Planes R, Serrero M, Hamimed I, Zurita E, et al. Trimeric heptad repeat synthetic peptides HR1 and HR2 efficiently inhibit HIV-1 entry. Biosci Rep. 2019;39(9).

31. Weng Y, Weiss CD. Mutational analysis of residues in the coiled-coil domain of human immunodeficiency virus type 1 transmembrane protein gp41. Journal of virology. 1998;72(12):9676–82.

32. Chan DC, Kim PS. HIV entry and its inhibition. Cell. 1998;93(5):681–4.

33. Aiewsakun P, Katzourakis A. Marine origin of retroviruses in the early Palaeozoic Era. Nat Commun. 2017;8:13954.

34. Hayward A, Cornwallis CK, Jern P. Pan-vertebrate comparative genomics unmasks retrovirus macroevolution. Proc Natl Acad Sci U S A. 2015;112(2):464–9.

35. Hayward A, Grabherr M, Jern P. Broad-scale phylogenomics provides insights into retrovirus-host evolution. Proc Natl Acad Sci U S A. 2013;110(50):20146–51.

36. Cui J, Zhao W, Huang Z, Jarvis ED, Gilbert MT, Walker PJ, et al. Low frequency of paleoviral infiltration across the avian phylogeny. Genome Biol. 2014;15(12):539.

37. Stenglein MD, Sanders C, Kistler AL, Ruby JG, Franco JY, Reavill DR, et al. Identification, characterization, and in vitro culture of highly divergent arenaviruses from boa constrictors and annulated tree boas: candidate etiological agents for snake inclusion body disease. mBio. 2012;3(4):e00180–12.

38. Aswad A, Katzourakis A. Convergent capture of retroviral superantigens by mammalian herpesviruses. Nat Commun. 2015;6:8299.

39. Shi M, Lin XD, Tian JH, Chen LJ, Chen X, Li CX, et al. Redefining the invertebrate RNA virosphere. Nature. 2016;540(7634):539–43.

40. Chuong EB, Elde NC, Feschotte C. Regulatory activities of transposable elements: from conflicts to benefits. Nat Rev Genet. 2017;18(2):71–86.

41. Chuong EB, Elde NC, Feschotte C. Regulatory evolution of innate immunity through co-option of endogenous retroviruses. Science. 2016;351(6277):1083–7.

42. Feschotte C, Gilbert C. Endogenous viruses: insights into viral evolution and impact on host biology. Nat Rev Genet. 2012;13(4):283–96.

43. Lima-Junior DS, Krishnamurthy SR, Bouladoux N, Collins N, Han SJ, Chen EY, et al. Endogenous retroviruses promote homeostatic and inflammatory responses to the microbiota. Cell. 2021;184(14):3794–811.e19.

44. Basta HA, Cleveland SB, Clinton RA, Dimitrov AG, McClure MA. Evolution of teleost fish retroviruses: characterization of new retroviruses with cellular genes. J Virol. 2009;83(19):10152–62.

45. Zagon IS, McLaughlin PJ. Opioid growth factor and the treatment of human pancreatic cancer: a review. World J Gastroenterol. 2014;20(9):2218–23.

46. Altschul SF, Gish W, Miller W, Myers EW, Lipman DJ. Basic local alignment search tool. J Mol Biol. 1990;215(3):403–10.

47. Xu Z, Wang H. LTR_FINDER: an efficient tool for the prediction of full-length LTR retrotransposons. Nucleic acids research. 2007;35(Web Server issue):W265–8.

48. Ellinghaus D, Kurtz S, Willhoeft U. LTRharvest, an efficient and flexible software for de novo detection of LTR retrotransposons. BMC Bioinformatics. 2008;9:18.

49. Katoh K, Standley DM. MAFFT multiple sequence alignment software version 7: improvements in performance and usability. Molecular biology and evolution. 2013;30(4):772–80.

50. Lu S, Wang J, Chitsaz F, Derbyshire MK, Geer RC, Gonzales NR, et al. CDD/SPARCLE: the conserved domain database in 2020. Nucleic Acids Res. 2020;48(D1):D265–d8.

51. Quinlan AR, Hall IM. BEDTools: a flexible suite of utilities for comparing genomic features. Bioinformatics (Oxford, England). 2010;26(6):841–2.

52. Krzywinski M, Schein J, Birol I, Connors J, Gascoyne R, Horsman D, et al. Circos: an information aesthetic for comparative genomics. Genome Res. 2009;19(9):1639–45.

53. Capella-Gutiérrez S, Silla-Martínez JM, Gabaldón T. trimAl: a tool for automated alignment trimming in large-scale phylogenetic analyses. Bioinformatics (Oxford, England). 2009;25(15):1972–3.

54. Kumar S, Stecher G, Li M, Knyaz C, Tamura K. MEGA X: Molecular Evolutionary Genetics Analysis across Computing Platforms. Molecular biology and evolution. 2018;35(6):1547–9.

55. Abascal F, Zardoya R, Posada D. ProtTest: selection of best-fit models of protein evolution. Bioinformatics. 2005;21(9):2104–5.

56. Nguyen LT, Schmidt HA, von Haeseler A, Minh BQ. IQ-TREE: a fast and effective stochastic algorithm for estimating maximum-likelihood phylogenies. Molecular biology and evolution. 2015;32(1):268–74.

57. Li J, Yu H, Wang W, Fu C, Zhang W, Han F, et al. Genomic and transcriptomic insights into molecular basis of sexually dimorphic nuptial spines in Leptobrachium leishanense. Nat Commun. 2019;10(1):5551.

58. Kim D, Langmead B, Salzberg SL. HISAT: a fast spliced aligner with low memory requirements. Nat Methods. 2015;12(4):357–60.

